# Protein interaction analysis of *Plasmodium falciparum* circumsporozoite protein variants with human immunoproteins explains RTS,S vaccine efficacy in Ghana

**DOI:** 10.1101/2021.12.29.474460

**Authors:** Cheikh Cambel Dieng, Colby T. Ford, Anita Lerch, Dickson Donu, Jennifer Huynh, Kovidh Vegesna, Jun-tao Guo, Daniel A. Janies, Linda Amoah, Yaw Afrane, Eugenia Lo

**Author notes:** These authors contributed equally (Cheikh Cambel Dieng, Colby T. Ford).

## Abstract

**Background:** The world’s first malaria vaccine RTS,S provides only partial protection against *Plasmodium falciparum* infections. The explanation for such low efficacy is unclear. This study examined the associations of parasite genetic variations with binding affinity to human immunological proteins including human leukocyte antigen (HLA) and T cell receptors (TCR) involved in RTS,S-induced immune responses.

**Methods:** Multiplicity of infections was determined by amplicon deep sequencing of merozoite surface protein 1 (*PfMSP1*). Genetic variations in the C-terminal of circumsporozoite protein (*PfMSP1*) gene were examined across 88 samples of *P. falciparum* collected from high and low transmission settings of Ghana. Binding interactions of *PfMSP1* variants and HLA/TCR were analyzed using `NetChop` and `HADDOCK` predictions. Anti-CSP IgG levels were measured by ELISA in a subset of 10 samples.

**Findings:** High polyclonality was detected among *P. falciparum* infections. A total 27 CSP haplotypes were detected among samples. A significant correlation was detected between the CSP and MSP multiplicity of infection (MOI). No clear clustering of haplotypes was observed by geographic regions. The number of genetic differences in *PfCSP* between 3D7 and non-3D7 variants does not influence binding interactions to HLA/T cells nor anti-CSP IgG levels. Nevertheless, *PfCSP* peptide length significantly affects its molecular weight and binding affinity to the HLA.

**Interpretations:** The presence of multiple non-3D7 strains among *P. falciparum* infections in Ghana impact the effectiveness of RTS,S. Longer *PfCSP* peptides may elicit a stronger immune response and should be considered in future version RTS,S. The molecular mechanisms of RTS,S cell-mediated immune responses related to longer CSP peptides warrants further investigations.

## Introduction

*Plasmodium falciparum* remains the most dominant malaria parasite in Sub-Saharan Africa that accounts for almost all mortality in malaria (1). Despite the many interventions in place, malaria burden remains high over years and mortality is the highest in young children (2). RTS,S is a pre-erythrocytic vaccine that aims to trigger the human immune system to defend against *P. falciparum* malaria (3). RTS,S is designed based on a portion of the central Asn-Ala-Asn-Pro (NANP) amino acid repeats (B-cell epitopes) and the C-terminal (T cell epitopes) of the circumsporozoite protein (CSP) of the 3D7 lab strain citeRN13. The C-terminal, precisely the Th2R and Th3R regions, encode epitopes recognized by the CD8+ and CD4+ T cells (Figure 1). The Th2R and Th3R portions (RTS) fuse to the native hepatitis B surface antigen (S) and self-assemble into virus-like particles exposed on the surface of the CSP protein (4–7). Phase III trials conducted at 11 sites across seven African countries indicated a 36-55% efficacy against clinical malaria in 5 to 17 monthold children who received three primary doses of RTS,S plus a booster at 20 months (3, 8–12). The explanation for such low efficacy is unclear.

**Fig. 1.**
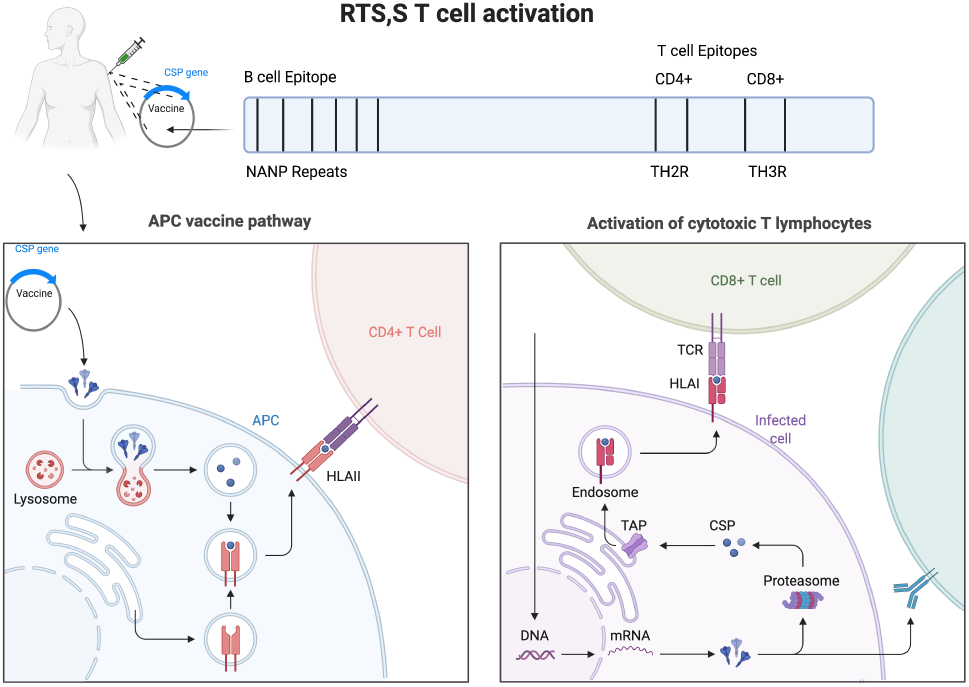
RTS,S T cell Activation.

High genetic diversity was reported in the *PfCSP* C-terminal as malaria transmission increases across geographic areas (13, 14). In high transmission areas, multiple genetically diverse parasite strains are common within the same host and in the community (15). Diverse parasite strains could harbor different CSP phenotypes contributing to variations in the invasion and or evasion mechanism of the parasites. Antibody-mediated immunity against blood-stage *P. falciparum* was strain-specific (16, 17). Functional responses elicited against a particular parasite strain do not yield comparable levels of inhibition against heterologous strains (18, 19). RTS,S efficacy against non-3D7 parasite strains was lower than that against 3D7 strain (20), further suggesting that allele-specific immunity is important in eliciting protection. Vaccine efficacy reduced from 50.3% for parasites with a perfect match to *PfCSP* 3D7 C-terminal sequence to 33.4% for parasites with any amino acid mismatch. Besides, vaccine efficacy declined as the number of amino acid differences increased (20). Mutational changes in amino acid sequence could alter CSP protein conformation and binding reaction of the CSP peptide with human leukocyte antigens (HLA), which is vital to the induction of T cell immune response, such as production and secretion of cytokines and the capacity to mediate cytotoxicity (21, 22). The recognition of *P. falciparum* by CD8+ and CD4+ T cell requires the intervention of antigen presenting cells (APCs) that display polypeptide fragments of *PfCSP* on their surfaces (Figure 1). The interaction between T cell and the CSP peptide is mediated by the HLA, which is expressed at the surface of APCs (23). The antigenic peptide recognition depends on the degradation of proteins by the 26S proteasome or lysosomal proteolysis, which generates appropriate peptide size to dock into the groove of HLA molecules (24–27). It is not clear if non-3D7 *PfCSP* variants exhibit different binding affinities to HLA and impact T cell recognition. Computational analyses using a machine learning-based model to predict protein structure, folding (28), macromolecules interactions and/or binding affinities have been the forefront of protein-ligand complexes studies (29). Such mechanistic insights would shed light on low RTS,S efficacy.

In Ghana, malaria accounts for 30% of outpatient attendances and 23% inpatient admissions in 2017(30). The country is characterized by heterogeneous transmission patterns driven by different climate and landscape features. Transmission is seasonal in the northern savannah, and perennial in the central forest and southern coastal areas (2). Ghana is one of the three African countries where RTS,S is being implemented along with Kenya and Malawi (2, 11). In this study, the multiplicity of infections (MOI) and CSP diversity among *P. falciparum* samples from different transmission settings of Ghana were compared. We aim to (1) determine the within and between host genetic diversity of *P. falciparum* across transmission settings; (2) clarify the effect of polyclonal infections on vaccine efficacy; (3) use protein predicted structure to identify mutational changes that may alter the overall conformation of the CSP protein and the binding affinity to HLA/T cell receptors. Our results have important implications on the development of next generation RTS,S. Namely, that CSP sequences that result in peptides of specific lengths may contribute to an increase in binding affinity between the HLA and T cell receptors.

## Methods

### Study sites and sample collection

Five sites were selected from three ecological zones of Ghana (31) including Pagaza (PZ) in Tamale Municipality and Kpalsogou (KG) in Kumbungu district in the northern savannah region; Duase (KD) in Konongo and Seikwa district in the central forest region; and Ada (AD) and Dodowa (DO) in the southern coastal region. Sample collection was conducted during June-July of 2019. About 200 *µ*L capillary blood samples were collected from 88 subjects who were diagnosed as malaria-positive by Access Bio CareStart^*TM*^ malaria rapid diagnostic test (RDT). DNA was extracted using Quick-DNA kit (Zymo Research) following the manufacturer’s protocol and stored in -20°C. *Plasmodium* species identification and DNA quantification was performed by real-time quantitative PCR of the 18S rRNA gene based on the published protocols (31).

### Amplicon deep sequencing and analysis

The 395-bp *PfMSP1* and the 321-bp C-terminal of *PfCSP* containing the Th2R and Th3R regions were amplified using previously published primers (32, 33). The Illumina partial adapters (34) were added to the purified PCR products prior to library constructions (See Table 1). Multiple samples were pooled and paired-end sequenced on the Illumina Hiseq 2500 (35). Coverage of at least 50,000 reads were obtained for each sample amplicon (sequences and sample information are available on NCBI under BioProject PRJNA783000).

**Table 1.**
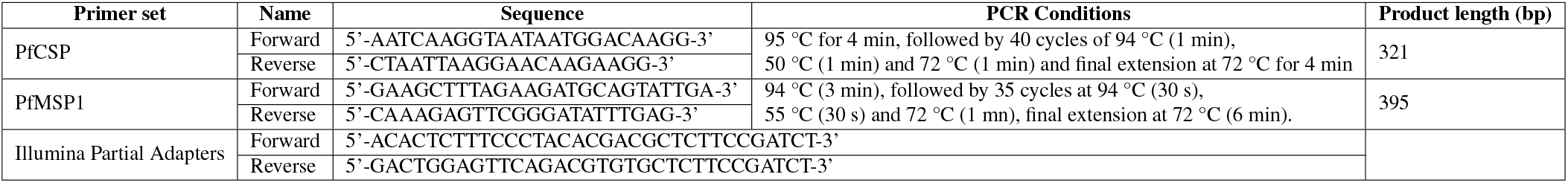
Sequences of the primers used in the study and corresponding references.

The `HaplotypR` package was used for SNP and haplotype detection based on *PfCSP* and *PfMSP1* sequences (36). Given that *PfMSP1* is length polymorphic, haplotypes were generated based on the size of the amplicon (Table 1), which were determined by iterative testing of various thresholds to distinguish between real haplotypes and random noise. MOI was calculated as the number of genetically distinct parasite clones co-infecting a single host with cut-off settings for haplotype calling requiring a haplotype to be supported by ≥ 3 reads in ≥2 samples (including independent replicates of the same sample) (36). A haplotype network was generated with the TCS for multiple sequence alignment evaluation and phylogenetic reconstruction (37). The construction of the haplotype network involves extracting the sequences two by two to build a `T-Coffee` library. The library was then used to obtain a Transitive Consistency Score (TCS) of every pair of aligned residues. Then, by averaging the pair-wise TCS score over all pairs of aligned residues in each target, SequenceTCS and the AlignmentTCS metrics were derived (38). The network was visualized using a TCS web-based program `tcsBU` (39).

### CSP peptide prediction and protein interaction modeling

To predict the resulting polypeptide expressed after proteolytic degradation of the CSP protein, the complete *PfCSP* haplotype sequences were scored using `NetChop 3.0` (40). `NetChop` provides a prediction score for each residue as to its probability of being expressed in a short peptide (~ 8 amino acids in length) yielded from the proteasomal degradation process. Residues with an expression probability ≥ 50% in the Th2R and Th3R regions of the protein were selected. These short peptide sequences were then assembled as 3D structures using the `Peptide Builder` package in Biopython (41). A PDB format file of each peptide was produced and used in subsequent interaction modeling. For comparison, we analysed the sequences through `netMHC 1` and `2` for TH2R and TH3R, respectively, to provide binding affinity predictions by position in the region. In `netMHC`, peptides are predicted if they were among the highest prediction score for HLA class I affinity, TAP transport efficiency, and C-terminal proteasomal cleavage as a weighted sum of the three individual prediction scores at the set threshold (42).

The interaction between CSP, HLA, and T cell receptor proteins is complex in that it requires the intervention of different components of the immune system (Figure 1). In humans, there are many different alleles of HLA-I and HLA-II molecules. However, these variants are expressed in distinct frequencies in different ethnic groups (43). According to the Allele Frequency Net Database, the HLA locus A (HLA-A) and HLA-DRB1 are the most frequent in Ghana and confer protection against severe malaria (44, 45).

Using the generated CSP peptide structure along with reference structures for the Human Leukocyte Antigen 2A (HLA-2A) and T Cell Receptor (TCR), we predicted the interaction of these three structures using `HADDOCK 2.4` (29). Specifically, we used PDB: 6TRN (chain A) for the HLA-2A structure and PDB: 6PY2 (chains D and E) for the TCR structure, renumbering residues according to `HADDOCK`’s requirements. We specified the interacting residues between the HLA structure, the CSP peptide, and the TCR structure by first docking the CSP peptide in the protein-binding groove of the HLA specifically between the *α*1 and *β*1 subunits (46). This HLA-CSP peptide complex was then docked at the antigen-binding site of the TCR on top of the *α* and *β* loci (46). The prediction of these interactions was derived from various biophysical factors including van der Waals energy, electrostatic energy, desolvation energy, and restraints violation energy, which were collectively used to derive a HADDOCK score to quantify changes in protein-protein interaction resulting from CSP peptide variations (47, 48).

### Measurement of anti-CSP antibody levels

Plasma from participant blood samples were examined for antibody titers against *PfCSP* by standardized enzyme-linked immunosorbent assays (ELISA) following a previous protocol (49). Recombinant IgG (BP055) was used as a standard calibrator for total IgG measurements. The *PfCSP* antigen was diluted to a concentration of 1 *µ*g/ml in carbonate buffer (pH 9.0) and coated at 100 µl/well onto ELISA plates at 4oC overnight, followed by washing with 1×PBS/Tween20 for four times. Plasma from malaria naïve individuals from Denmark was used as negative control samples. Absorbance was measured using the ELx808 Absorbance Reader (Biotek, USA) and resulting optical density values were converted to IgG concentration using ADAMSEL (Ed Remarque, BPRC). This process workflow is depicted in Figure 2.

**Fig. 2.**
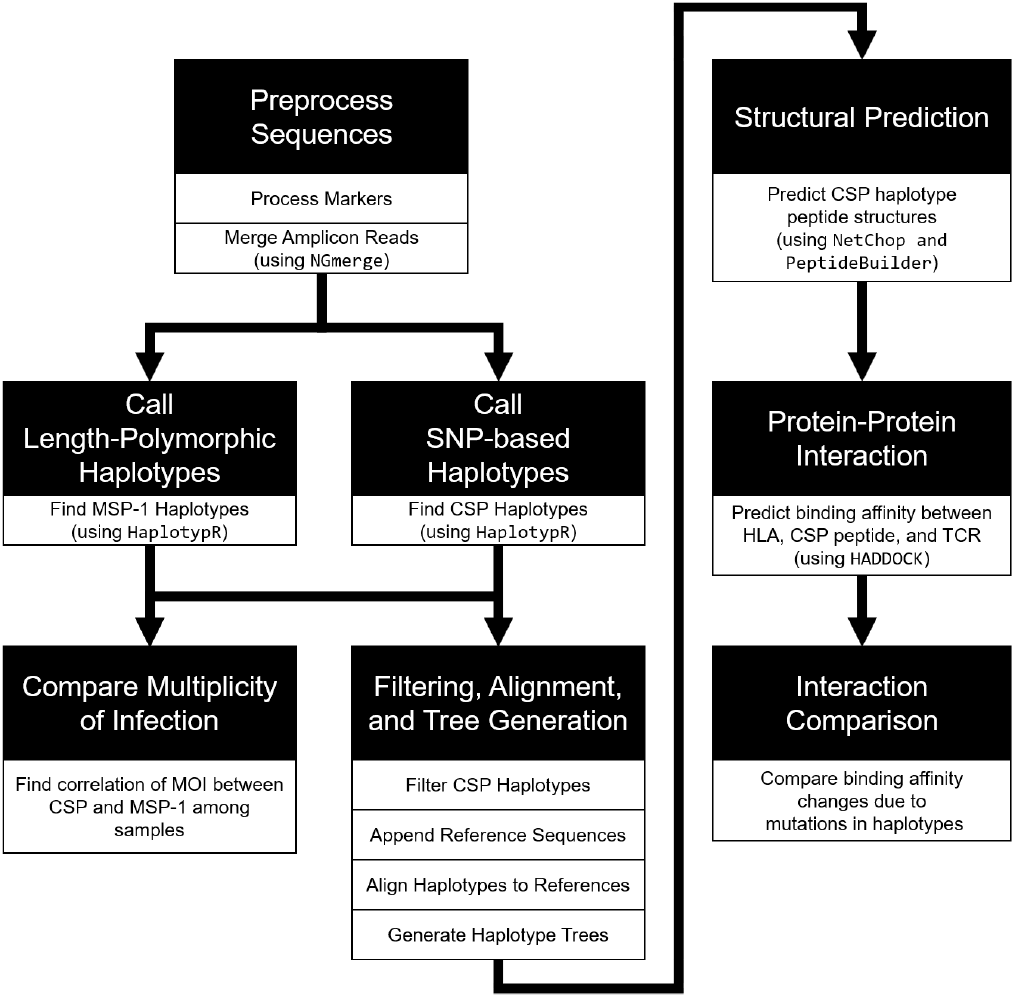
Process workflow for generating and comparing CSP haplotypes and assessing human immunoprotein interaction.

## Results

### Multiplicity of infection (MOI) and Sequence Comparison

Based on *PfMSP1*, the proportion of polyclonal samples among the study sites was 76%, with the mean MOI of 2.63 among the 88 samples (Table 3). A total of 30 haplotypes were detected, ranging from 1 to 6 distinct clones within a single host (Figure 3). Considerable size polymorphisms were observed among different the *PfMSP1* haplotypes. Polyclonal infections were detected in 80% of the samples from the North and all samples in the central and southern regions. The mean MOI was highest in central followed by the north and south Ghana, but such difference was not significant(Table 3)

**Table 2.** `HaplotypR` settings for generating CSP and MSP1 haplotypes.

| Setting | CSP | MSP1 |
| --- | --- | --- |
| Method | SNP-based | Length Polymorphic |
| Minimum Mismatch Rate (for SNPs) | 0.50 | NA |
| Minimum Coverage | 3 | 3 |
| Detection Limit | 1/50 | 1/100 |
| Minimum Occurrence | 3 | 2 |
| Minimum Coverage per Sample | 25 | 25 |

**Table 3.** Comparison of multiplicity of infections (MOI) and percentage of polyclonal samples based on *PfMSP1*, as well as nucleotide (Pi) and haplotype (Hd) diversity based on *PfCSP* among samples from north, central, and south Ghana. (*Note: This only represents 83 of the 88 samples where we have both CSP and MSP-1 sequences*.)

|  |  | *PfMSP1* |  | *PfCSP* |  |
| --- | --- | --- | --- | --- | --- |
|  | Total Samples | % Polyclonality | Multiplicity of Infections (mean, [range]) | Nucleotide Diversity | Haplotype Diversity (Hd) |
| **North** | 27 | 80 | 2.9, [1-6] | 0.017 | 0.951<br>(-0.011) |
| **Central** | 27 | 100 | 3.1, [1-6] | 0.017 | 0.945<br>(-0.009) |
| **South** | 29 | 100 | 2, [0-4] | 0.018 | 0.938<br>(-0.009) |

**Fig. 3.**
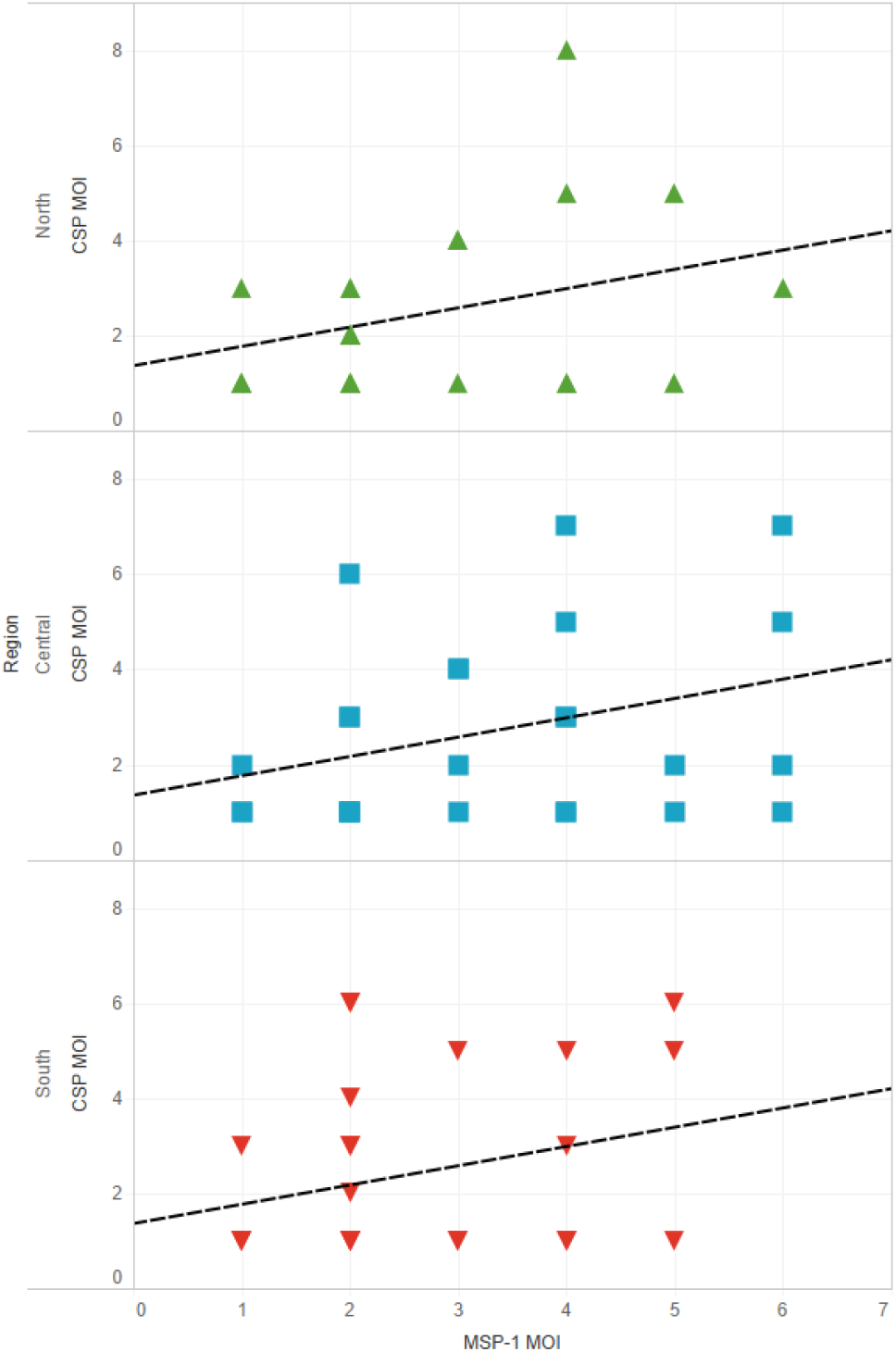
Relationship of CSP and MSP1 minimum multiplicity of infection by geographic region. *(Overall R: 0*.*3197, p-Value: 0*.*0052)*

For *PfCSP*, 27 haplotypes were detected among samples, ranging from 1 to 8 haplotypes/clones within a host (Figure 3). Nucleotide diversity (Pi) and haplotype diversity (Hd) were similar among the three studied regions (Table 3). Though the Pi values were found to be less than 0.02 in each region, Hd values ranged from 0.938–0.951, indicating a high clonal diversity in the parasite populations. For all samples, MOI values estimated from the *PfMSP1* and *PfCSP* were significantly correlated with one another (R = 0.33; p-value = 0.005). When samples were analyzed by regions, only those in the north showed a significant correlation (R = 0.41; p-value = 0.039). The northern, central, and southern populations shared similar *PfCSP* haplotypes, and no clear clustering of haplotypes was observed by geographic regions. Of the 27 *PfCSP* haplotypes, haplotype IDs CSP-10 and CSP-31 were most closely related to the 3D7 haplotype with only one SNP difference (Figure 4). By contrast, haplotypes with IDs CSP-15, CSP-117, and CSP-22 were the most distant from 3D7, each with 11-12 nucleotide differences. For CSP-117, mutations observed at the Th2R and Th3R regions were non synonymous and thus, this haplotype has identical amino acid sequence as 3D7. (Figure 5)

**Fig. 4.**
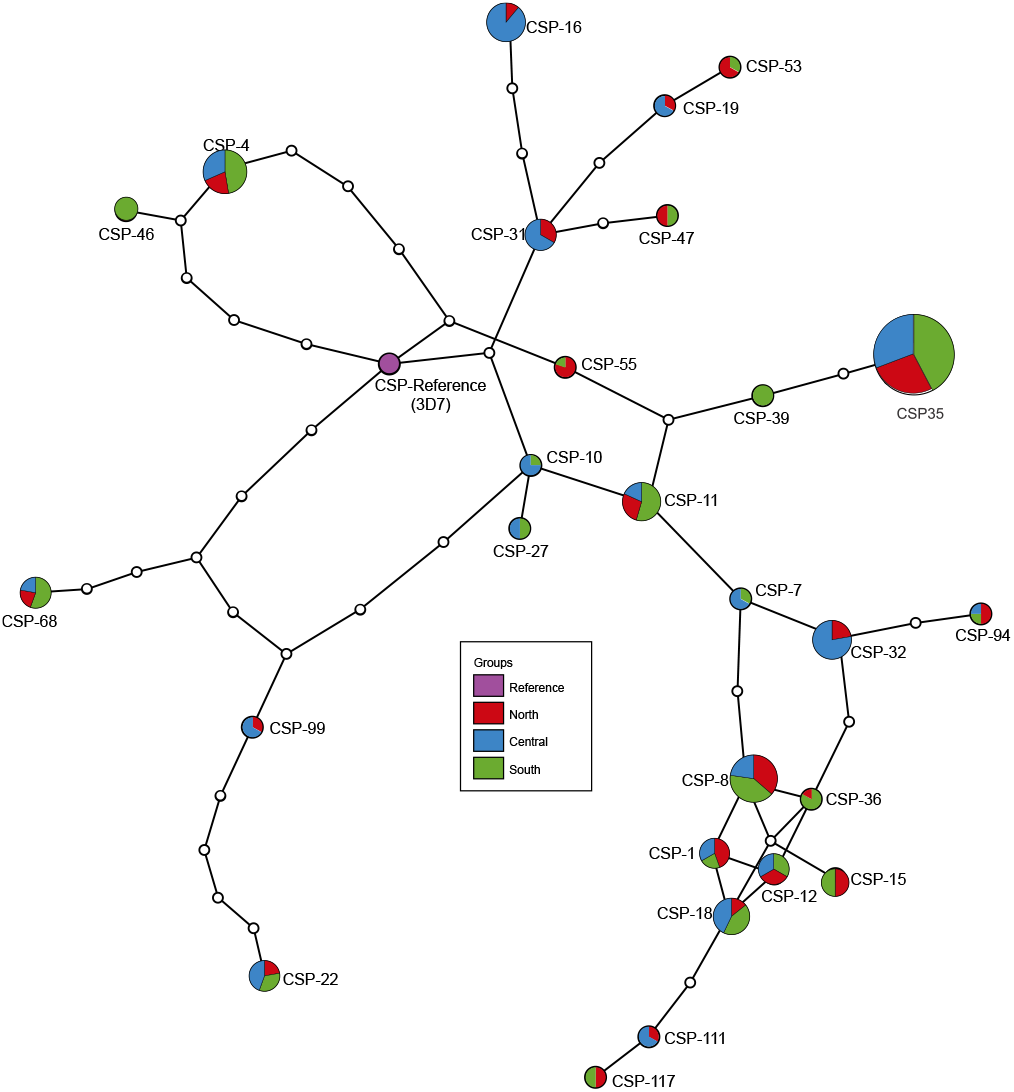
Haplotype network of the 27 CSP haplotypes. This was generated using `TCS` (50) and then colorized with region information using `tcsBU` (51).

**Fig. 5.**
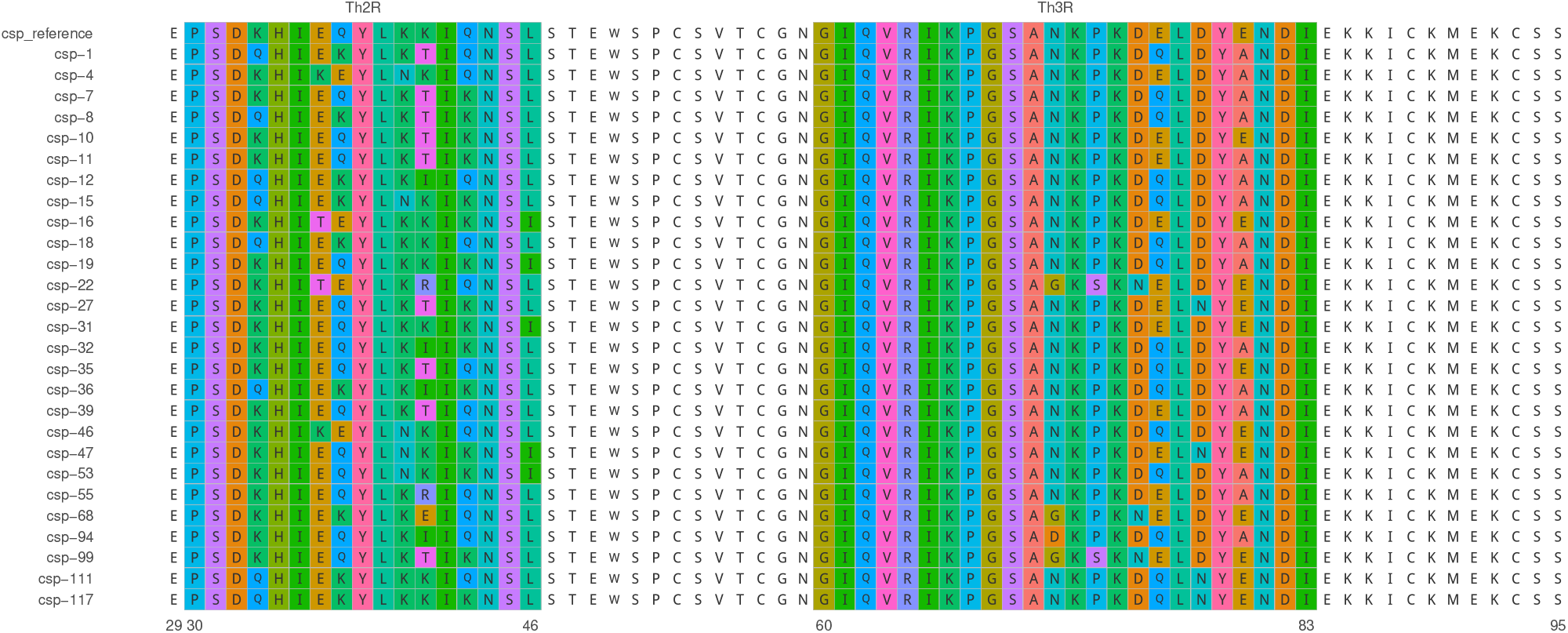
Sequences of the 27 CSP haplotypes found in this study. The Th2R and Th3R regions are highlighted to show variations from the 3D7 reference sequence.

### Binding affinity of *PfCSP* variants

The C-terminal of the *PfCSP* sequences that contained the Th2R and Th3R regions (known as T cell epitope regions (52)) were analyzed separately for binding affinity. In the Th3R region, four peptides (VIELYE, VIQLYA, VIELYA, VIQLYE) were detected based on six amino acid residues (Figures in Table 8). HAD-DOCK scores of these peptides with the HLA-TCR complex were very similar without clear differences (Table 4).

**Table 4.**
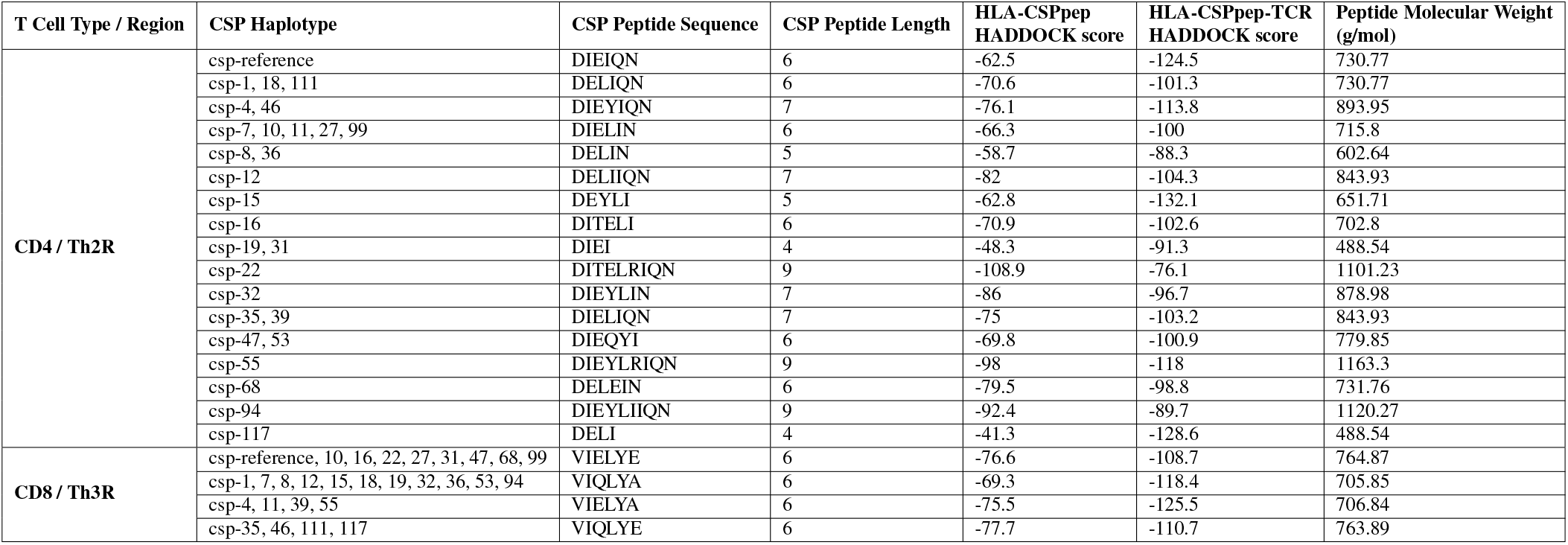
Summary of the interaction of the different CSP peptides and both HLA molecules and T cell structures.

Compared to Th3R, the Th2R region of *PfCSP* was more variable and contained 17 distinct peptide sequences (Figures in Table 7). The binding peptides of Th2R predicted by `NetChop` ranged from 4 to 9 residues in length and all begin with a glutamate (D) residue. As shown in Figure 6, there is a significant correlation between a peptide’s molecular weight and the interaction between the HLA and CSP peptide. Longer peptides have a better binding affinity within the HLA groove (R = -0.9415; p-value = <0.0001).

**Fig. 6.**
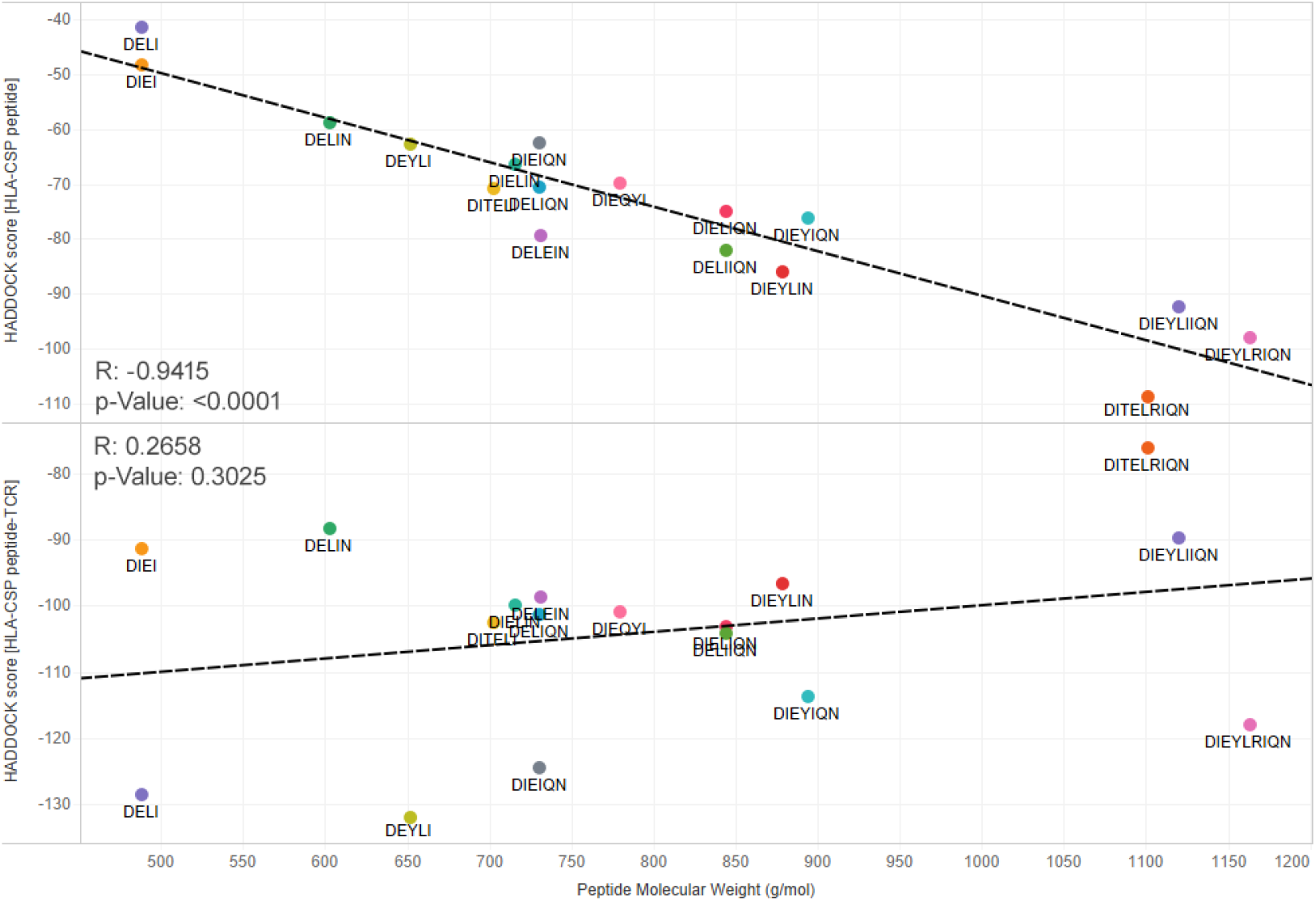
Correlation plots of HADDOCK scores by Th2R CSP haplotype peptide molecular weight.

However, these trends were not observed when we included the T cell structure to the to the HLA-CSP peptide (R = 0.2658; p-value = 0.3025). The HADDOCK scores of Th2R peptides were compared among haplotypes that were closely related and distant from the 3D7 reference. For instance, CSP-15 one of the most distant haplotypes, that provides a 5-residue peptide, showed the lowest HADDOCK score of -132.1 (the highest binding affinity; 6.1% higher than 3D7). Similarly, CSP-117, which provides a 4-residue peptide, showed a HADDOCK score of -128.6 (3.3% better than 3D7), the second-best interaction with the HLA and T cell complex. By contrast, of all haplotypes, CSP-22 and CSP-8 showed the lowest binding affinity. CSP-22, which provides a 9-residue peptide. showed a HADDOCK score of -76.1 (the lowest binding affinity; 38.9% less than 3D7). CSP-8, which provides a 5-residue peptide, showed a HADDOCK score of -80.3 (35.5% less than 3D7). While CSP-10 and CSP-31 were the two most closely related haplotypes to 3D7, their HADDOCK scores were markedly different. CSP-10, which provides a 6-residue peptide (DIELIN), has a similar HADDOCK score as 3D7, but CSP-31, which contains a 4-residue peptide (DIEI), has one of the lowest HADDOCK scores. See Table 4. These findings indicated that the number of genetic variations from 3D7 does not correlate with the binding affinity with the HLA and T cell complex.

To further elucidate if residues with larger side chains, such as tyrosine or glutamine, may cause the CSP peptide to block the HLA groove, thereby reducing the binding interaction, we compared the HADDOCK scores and other evaluation metrics for peptides containing and lacking tyrosine or glutamine. The *χ*^2^-statistic indicated no significant difference among these peptides. (Table 5; Figure 7)

**Table 5.** Kruskal-Wallis test results indicating the chi-square and p-values of binding affinity matrices between CSP peptides that contain or lack tyrosine or glutamine. *No significant correlations below the α = 0*.*05 level were found*.

| HLA-CSP peptide-TCR Metric | Peptide Contains | $\chi^2$ Statistic | p-value |
| --- | --- | --- | --- |
| Restraints violation energy | Glutamine | 2.676 | 0.102 |
| Restraints violation energy | Tyrosine | 0.091 | 0.763 |
| Desolvation energy | Glutamine | 3.177 | 0.075 |
| Desolvation energy | Tyrosine | 0.041 | 0.840 |
| Electrostatic energy | Glutamine | 1.815 | 0.178 |
| Electrostatic energy | Tyrosine | 1.222 | 0.269 |
| van der Waals energy | Glutamine | 1.335 | 0.248 |
| van der Waals energy | Tyrosine | 0.205 | 0.650 |
| HADDOCK score | Glutamine | 0.231 | 0.630 |
| HADDOCK score | Tyrosine | 0.364 | 0.546 |

**Fig. 7.**
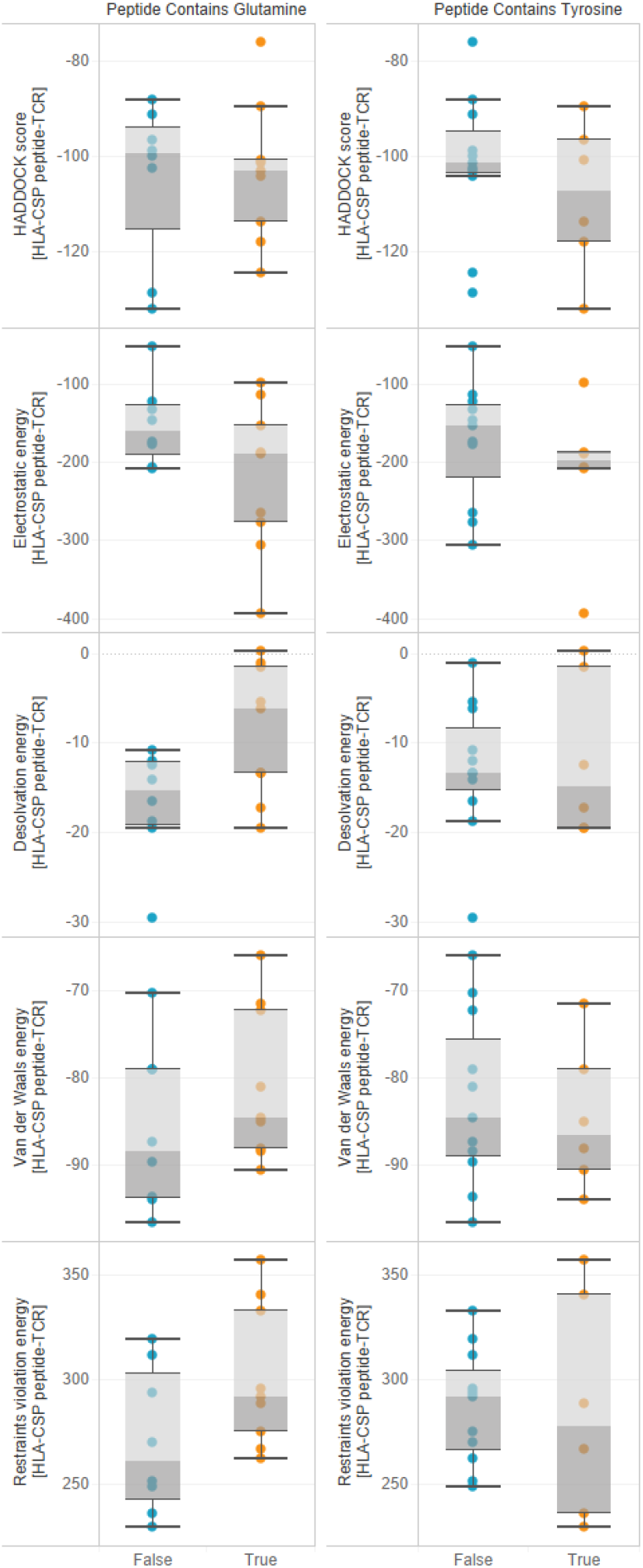
Boxplots breakdown of statistics of Th2R CSP peptides by the presence or absence of glutamine or tyrosine.

### Association of peptide length with binding affinity

To determine if peptide molecular weight (based on peptide length) of CSP influences the binding affinity of HLA and T cell complex, we created correlation plots between HAD-DOCK score and various peptide molecular weight derived from lysosomal degradation (Figure 6). A strong negative correlation was detected between the HADDOCK score of the HLA-CSP peptide and the molecular weight of the CSP peptide (R = -0.94; p-value = <0.001), suggesting that longer CSP peptides confer to stronger binding affinity. However, when TCR was added to the HLA-CSP complex, the HAD-DOCK score was not significantly correlated with the molecular weight of the complex (R = 0.2658; p-value = 0.3025). These findings suggest that longer peptides may bind better within the HLA groove and the size of the *PfCSP* peptide predicted based on proteolysis may play role in recognition by T cell receptors. Other binding metrics were also assessed in Table 6.

**Table 6.**
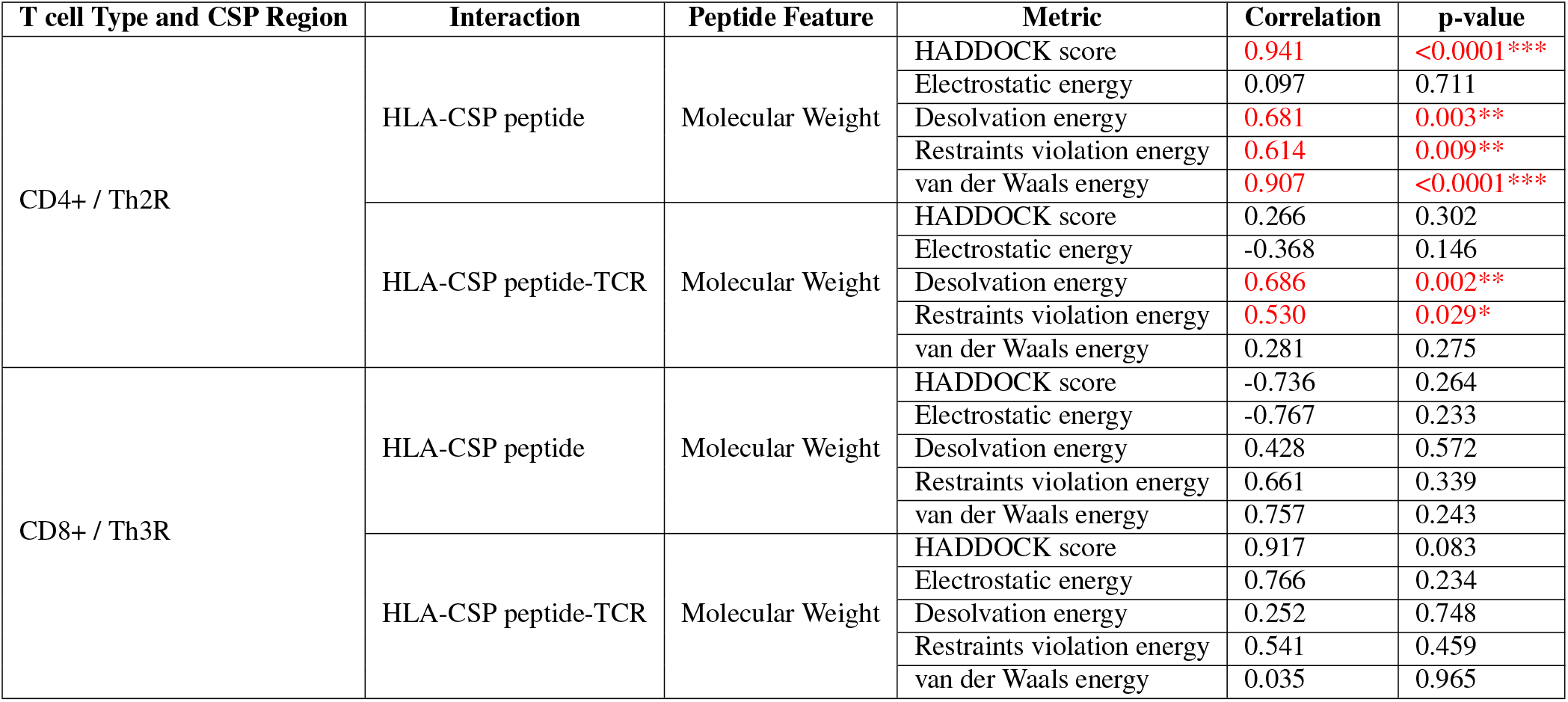
Correlations of peptide features against docking metrics by interaction and T cell type and *PfCSP* region. *Significant correlations below the α = 0*.*05 level are denoted in red and with one or more asterisks (*p<0*.*05; **p<0*.*01; ***p<0*.*001)*.

**Table 7.**
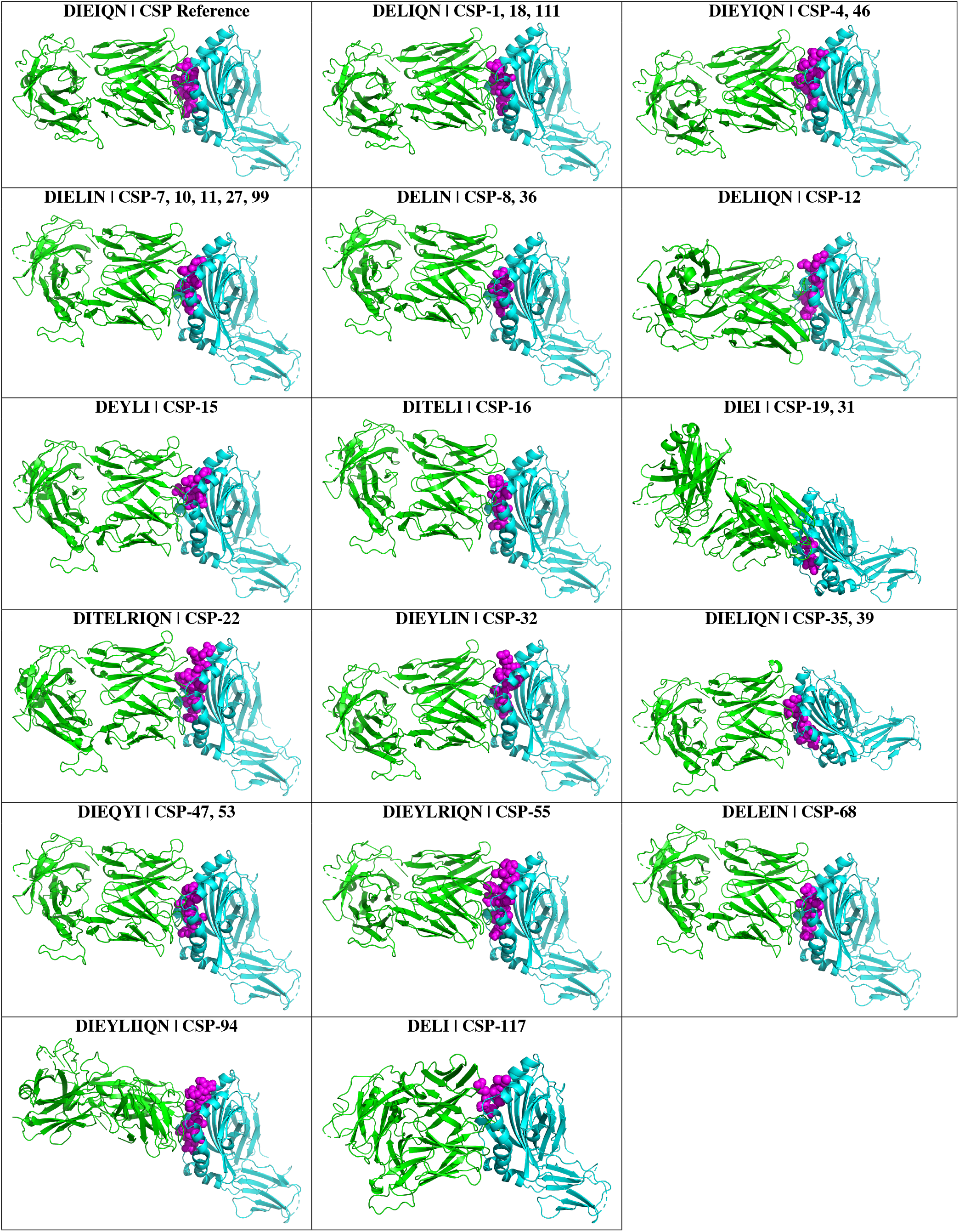
Th2R Region HADDOCK Protein Interaction Predictions using MHC Class II HLA-DR (PDB: 6V1A) and T cell receptor T594 (PDB: 6PY2). *Note that the TCR structure is shown in green, the CSP peptide is shown in magenta, and the HLA structure is shown in cyan*.

**Table 8.**
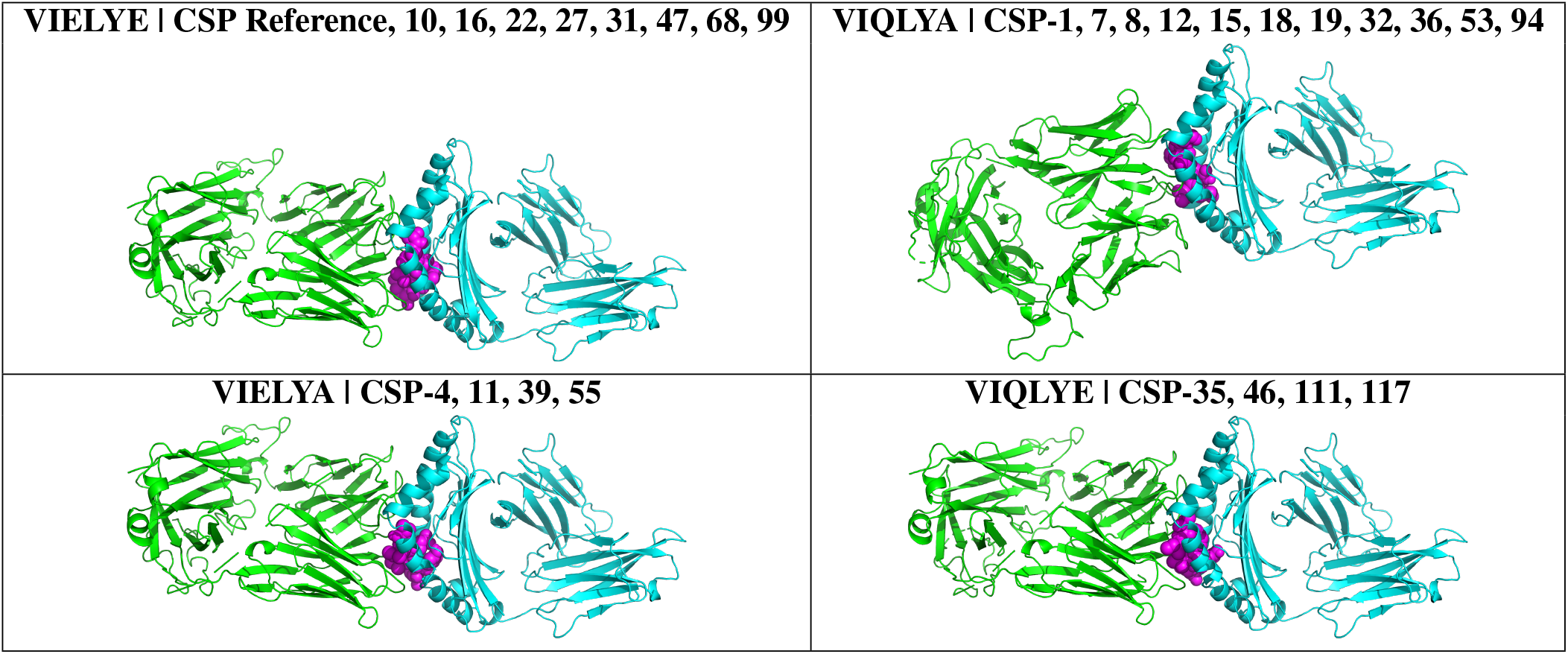
Th3R Region HADDOCK Protein Interaction Predictions using MHC Class I HLA-A2*01 (PDB: 6TRN) and T cell receptor T594 (PDB: 6PY2). *Note that the TCR structure is shown in green, the CSP peptide is shown in magenta, and the HLA structure is shown in cyan*.

### Association of *PfCSP* antibody levels with ligand-peptide binding affinity

For a subset of samples, *PfCSP* antibody levels were measured to determine if a binding affinity correlation exists between CSP peptide and HLA/TCR complex. No significant correlation was found between *PfCSP* antibody level and the binding affinity of CSP peptide with HLA-CSP (R = 0.61; p-value = 0.059) nor HLA-CSP TCR complex (R = 0.23; p-value = 0.52). The antibody level was also not associated with qPCR-based parasitemia among the infected samples (R = 0.58; p-value = 0.082; Figure 8)

**Fig. 8.**
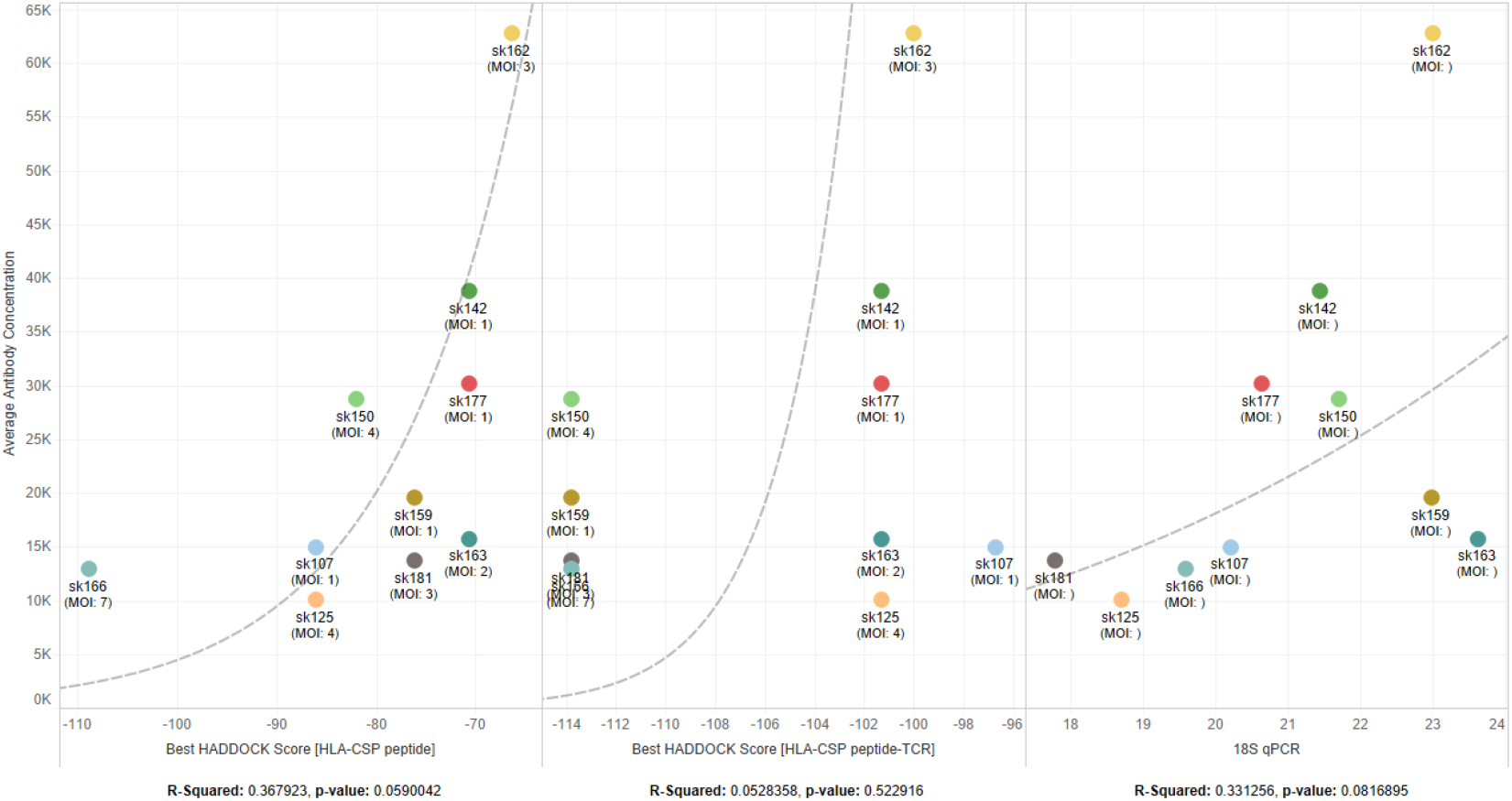
Correlation plots showing a potential, though not significant, logarithmic relationship between antibody concentration and binding affinity.

## Discussion

*Plasmodium falciparum* infections in highly endemic areas are usually complex and polyclonal, which explains the great genetic polymorphism within the speciescite (53). Such diversity could impact different antimalarial interventions such as the emergence of resistant parasite strains and break-through infections of malaria vaccines. Here, we predicted that the presence of multi-clone infections and high prevalence of the non-3D7 *PfCSP* variants may allow the parasites to escape RTS,S-induced immunity and explain the relatively low efficacy observed in phase 3 trial (9). Consistent with other studies, *PfMSP1* is more variable than *PfCSP* in distinguishing parasite clones within hosts (36). The positive correlation of *PfCSP* MOI and *PfMSP1* MOI values suggested that individuals who were infected with a higher number of parasite clones harbored multiple *PfCSP* variants that could impact immune responses induced by RTS,S. Among the Ghanaian samples, the genetic heterogeneity of the *PfCSP* was relatively high, with up to six distinct haplotypes within a sample. While three of the samples had identical Th3R sequences as the 3D7 reference, no samples had the 3D7-type Th2R sequences. This result agrees with a previous study that found no polymorphism in Th3R, suggesting that this region may be under balancing selection (22, 54). The four Th3R peptides (VIELYE, VIQLYA, VIELYA, VIQLYE) had similar HADDOCK scores and conceivably similar binding affinity.

In this study, a clear correlation was detected between the molecular weight of CSP peptides and binding affinity when the peptide was docked in the protein-binding groove between the *α*1- and *β*1-subunits of HLA. However, when the HLA-CSP complex was docked at the antigen-binding site of the TCR structure, the molecular weight of the peptide was not significantly associated to binding affinity. It is possible that longer peptides with exposed side chains allow for a better interaction with the HLA receptor, but the size of the entire CSP-HLA complex does not impact its interaction with T cell receptors (26, 55). The peptide-binding groove of HLA-I is known to contain deep binding pockets surrounded by polymorphic sidechains, which imposes tight constraints on the residues and overall peptide length to achieve binding. By contrast, the binding groove of HLA-II is open, allowing peptides of various lengths to fit into the binding groove (56). This explains the challenge in predicting interactions of the HLA-II-peptide compared to HLA-I-peptide complexes. Peptide length has been shown to affect the interactions with HLA molecules. For instance, a study of CD8+ T cell response to the BZLF1 protein of the Epstein-Barr virus showed that individuals with HLA-B*18:01 responded strongly and exclusively to an octamer peptide and those with HLA-B*44:03+ responded to a dodecamer peptide (57), highlighting the importance of peptide length for T cell response (58).

Previous studies have shown that vaccine efficacy declined as the number of amino acid differences increased, suggesting that allele-specific immunity is important in eliciting protection (20). In Ghana, multiple CSP haplotypes are present within a polyclonal sample and most CSP haplotypes are markedly different from the 3D7 strain at the Th2R region. Haplotypes that were most closely related to 3D7 could have peptide length and/or and molecular weight different from 3D7 after proteolysis and exhibit completely different binding affinity with HLA. This finding suggests that the CSP peptide length rather than amino acid differences alone determine binding affinity and downstream immune responses. Further, residues with larger side chains such as tyrosine or glutamine in the CSP peptide have been shown to block the HLA groove (59), thereby reducing binding interaction. Nevertheless, our analyses showed no significant effect of tyrosine or glutamine on the docking of the CSP peptides.

Our results indicated no significant correlation between anti-CSP antibodies and binding affinity for both HLA-CSP and HLA-CSP-TCR complexes. However, only the most prevalent haplotype within the sample was investigated and the presence of other variants may affect antibody production. A recent study showed that high proportions of strain-specific antibody responses are likely to be elicited during high transmission season due to increased MOI and in turn, a higher preponderance of cross-reactive antibodies (18).

### Limitations

It is important to note that this study has a few limitations. First, our samples may represent only a portion of CSP variants present in Africa. None of the haplotypes in our samples were similar to the 3D7 reference. Including samples with a high prevalence of 3D7-matching haplotypes at the Th2R and Th3R regions would allow us to compare docking results at the T cell epitopes and vaccine efficacy with samples of mostly non-3D7 haplotypes as in the present study. RTS,S immune responses induced by other variants merits further investigations. Second, the HLA-2A and HLA-DR1 were selected in the protein modeling analysis by their high prevalence in Ghana and potential protection against severe malaria (60), even though there are several classes of HLA-I and HLA-II molecules in other ethnic groups that may exhibit different binding affinities (61). Similarly, the choice of the T cell receptor T594 was limited by the lack of representative T cell receptor structures bound to a *Plasmodium* protein in the Protein Data Bank. Third, only Th2R and Th3R regions of PfCSP were examined but epitopes in the N-terminal and NANP repeat regions may also affect binding interactions with HLA/TCR and corresponding immune responses. Future studies should validate the *in silico*-derived candidate sequences *in vitro* using custom-designed assays to investigate the HLA and TCR interactions. Also, the impact of *PfCSP* diversity in the N-terminal and central repeat regions on B-epitope recognition and vaccine efficacy remains unclear.

## Conclusions

This study reveals a high level of polyclonality in *P. falciparum* infections across different transmission settings of Ghana. Mutations were found predominantly in the Th2R region of *PfCSP* and the most prevalent *PfCSP* haplotypes are not closely related to the 3D7 reference. The length of the Th2R peptide is significantly correlated with binding affinity with the HLA molecules but genetic relatedness to 3D7 does not impact binding interactions. These findings provide new insights into low vaccine efficacy and inform development of next generation RTS,S.

## Future Work

In our upcoming research, we plan to validate the *in silico*-derived candidate sequences *in vitro* using custom-designed assays to test HLA and TCR interaction. Similar analyses are needed to further investigate the binding interactions and impact of other potential CSP haplotypes on vaccine design in other parts of Africa and the world using our methodologies described in this work.

## Supplementary Materials

All *PfMSP1* and *PfCSP* amplicon reads where uploaded to the National Center for Biotechnology Information Short Read Archive (NCBI SRA) under BioProject number: PRJNA783000.

All data, scripts, and results from this work are available at GitHub.com/colbyford/Ghana_CSP_Haplotypes.

## ACKNOWLEDGEMENTS

We thank the field team from The University of Ghana for their technical assistance in sample collection; the communities and hospitals, for their support and willingness to participate in this research; and undergraduates at The University of North Carolina Charlotte who were involved in sample processing.

## FUNDING

This research was funded by the 2019 University of North Carolina at Charlotte Faculty Research Grant. We also acknowledge funding and logistical support from several entities of the University of the North Carolina at Charlotte including: The Department of Bioinformatics and Genomics, the Bioinformatics Research Center, University Research Computing, the College of Computing and Informatics and the College of Liberal Arts and Sciences. We are grateful for funding from the Belk Family.

## COMPETING FINANCIAL INTERESTS

The authors declare no conflicts of interest.

## Notes

https://github.com/colbyford/Ghana_CSP_Haplotypes

https://www.ncbi.nlm.nih.gov/bioproject/PRJNA783000/

### Competing Interest Statement

The authors have declared no competing interest.

